# Permethrin elicits chemoreceptive responses on different *Anopheles gambiae* sensory appendages

**DOI:** 10.1101/2025.09.05.674203

**Authors:** Sassan S. Kambou, Adeline Valente, Philip Agnew, Anna Cohuet, David Carrasco

**Affiliations:** MIVEGEC, Univ. Montpellier, IRD, CNRS, Montpellier, 34394, France.; Institut de Recherche en Sciences de la Santé (IRSS), Centre National de Recherche Scientifique et Technique (CNRST), Bobo-Dioulasso, 01 BP 545, Burkina Faso

**Keywords:** pyrethroid, insecticide detection, *Malaria mosquito*, tarsi, antennae, palpi

## Abstract

**BACKGROUND:** Non-contact detection of pyrethroid insecticides by malaria mosquitoes has been unveiled and may contribute to the evolution of mosquito behavioral modifications against vector control tools. However, the mechanisms underlying this detection are not yet fully understood. It has been hypothesized that the spatial repellency of pyrethroids may be mediated by chemosensory receptors and/or via the activation of voltage-gated sodium channels (VGSCs). This study aimed to explore these two hypotheses by identifying which chemosensory appendages in *Anopheles gambiae* are involved in the non-contact detection of permethrin, a widely used pyrethroid in malaria control.

**RESULTS:** Behavioral responses to permethrin headspace were recorded in female *An. gambiae*, in which specific sensory appendages were either removed or coated with resin to impair their chemosensory function. Additionally, electrophysiological recordings were performed on different sensory appendages: antennae, palpi and tarsi, to characterize their electrophysiological activity after permethrin stimulation. The behavioral assays revealed that tarsi were primarily responsible for mediating mosquito takeoff responses after permethrin headspace delivery. This finding was supported by significant electrophysiological tarsal responses to the insecticide. In contrast, removal of the antennae did not alter behavioral responses, although electroantennogram recordings indicated neural activity in response to permethrin. The palps showed neither behavioral nor electrophysiological responses.

**CONCLUSION:** These findings indicate that permethrin is detected through two distinct sensory appendages, tarsi and antennae, but with varying behavioral output. Such appendage-specific detection favors the hypothesis that permethrin detection and the associated behavioral output is mediated by chemosensory receptors rather than by VGSCs. Nonetheless, further investigations are needed to identify the chemosensory receptors and pathways involved in pyrethroid insecticide detection in malaria mosquitoes.

## 1 INTRODUCTION

*Anopheles gambiae* is one of the main vector species transmitting malaria parasites in sub-Saharan Africa. Vector control based on pyrethroid insecticides, such as insecticide treated nets (ITNs) and indoor residual spraying (IRS), still remains one of the most effective strategies to reduce malaria transmission (1,2). However, mosquitoes resistance to insecticides is globally increasing and jeopardizes the efficacy of vector control measures (3–6). Several non-exclusive mechanisms of genetic and physiological resistance have been identified for most of the major malaria vectors (7,8). In addition, behavioral resistance mechanisms that partially or completely reduce mosquito exposure to insecticides have also been described (9–12).

Various forms of insecticide-related behavioral adaptations have been hypothesized [see (11)]. Those enabling mosquitoes to avoid direct contact with an insecticide treated surface are expected to be the most advantageous as the deleterious effects of the toxin can be avoided. However, some of the proposed examples of avoidance behavior require mosquitoes to detect the presence of the insecticide before contact. In a recent behavioral study, we showed that *An. gambiae* female mosquitoes, with or without target-site mediated insecticide resistance, were able to detect without contact three of the most common insecticides used in vector control: permethrin, alpha-cypermethrin and deltamethrin (13). Moreover, recent studies with *Aedes* mosquitoes have similarly revealed that other pyrethroid molecules (i.e., transfluthrin and bioallethrin) and natural pyrethroid analogs (i.e., pyrethrins) exert a repellent effect at a distance (14–18). In these studies, two non-mutually exclusive functional mechanisms have been proposed to explain the non-contact repellency to pyrethroids. On one hand, pyrethrins and the pyrethroids tested have been shown to be agonists of olfactory receptors. This suggests the implication of the common insect olfactory pathway in the detection of these compounds, which could eventually trigger a repellent behavioral effect. On the other hand, pyrethrins and pyrethroids are also (hyper-) activators of voltage-gated sodium channels (VGSCs) (19), hence their insecticidal properties. VGSCs activation could also elicit the repellent behavioral effect of these insecticide molecules (15,18), independently of sensory receptors mediation. Although a considerable phylogenic distance exists between *Aedes* and *Anopheles* species, similar mechanisms behind the non-contact chemosensory detection of pyrethroids by *Anopheles sp.* mosquitoes could exist.

Nonetheless, all previous functional studies on non-contact detection of pyrethroids have focused on the antennae, i.e., the main olfactory appendage of insects. However, mosquito chemosensory appendages also include: the maxillary palps, the labial palps, internal surfaces of mouthparts, distal leg segments and wing margins. All these appendages are covered with hair-like or dome shaped cuticular structures called sensilla (20–23). Sensilla number and morphological type vary depending on the species and body location (24). Sensilla are innervated by the dendrites of one or more chemosensory neurons (25), which express sensory receptors in their membranes. There are three main receptor families involved in mosquito chemo sensation: odorant receptors (ORs), ionotropic receptors (IRs), and gustatory receptors (GRs) (21,26–28), and their expression varies depending on the appendage. In *Anopheles* mosquitoes, for instance, the most expressed chemosensory receptors in the antennae are ORs and IRs; GRs, ORs and IRs in the palps, and GRs and IRs in the tarsi (20,29–31). Therefore, the appendage-specific distribution of chemosensory receptors contrasts with the broad distribution of voltage-gated sodium channels (i.e., pyrethroids target site), which in insects are found on the membranes of most excitable cells across the entire nervous system. This suggests that an appendage-specific detection of these compounds would denote a detection mainly through chemoreceptors, whereas a non-specific response may be in favor of a detection mainly mediated by voltage-gated sodium channels.

In order to address these hypotheses, we performed a series of behavioral and electrophysiological experiments to investigate the role of the main *Anopheles gambiae* chemosensory appendages in the non-contact detection of a commonly used pyrethroid insecticide, permethrin.

## 2 METHODS

### 2.1 Mosquito strain

The insecticide-susceptible strain of *Anopheles gambiae s.s.* Kisumu was used for all experiments (32). Mosquitoes were reared at Vectopôle insectary at Institut de Recherche pour le Développement (IRD) in Montpellier (France) under standard insectary conditions. Mosquitoes were maintained at 27 °C ±1, 70-80% relative humidity and a 14h:10h L:D cycle throughout their development. Larvae were fed with fish food flakes (TetraMine Flakes, Tetra, Germany), whereas adults were provided with 10% of sucrose solution.

### 2.2 Chemical compounds

Permethrin, DEET, geraniol, acetophenone, 1-octen-3-ol and ammonia water were obtained from Sigma-Aldrich (details in Table S1, supplementary information). All compounds were diluted in 99.9% acetone (Sigma-Aldrich) at different concentrations depending on the experiment (see below for concentration details). Acetone was thus used alone as negative control. DEET, geraniol, 1-octen-3-ol and ammonia water were used as positive controls in behavioral assays and/or electrophysiology experiments.

### 2.3 Mosquito preparation

The main chemosensory appendages of the mosquitoes; antennae, maxillary palps, proboscis and tarsi were sequentially rendered nonfunctional to determine their individual role in permethrin chemosensory detection. For that, female mosquitoes were placed into paper cups covered with a mesh and temporarily immobilized by placing the cups in an ice box for 2 minutes.

Once immobilized, the mosquitoes were manipulated over a cold plate under a binocular microscope according to five different treatments; in treatments (i)-(iii), respectively, the antennae, palps or proboscis were excised at their base using micro-scissors. In treatment (iv), all three of the appendages in treatments i-iii were excised. For treatment (v), the tarsi were recovered with UV curing resin (ABS-like Photopolymer Resin, UV wavelength 405 nm) in order to occlude the sensory sensilla present without impeding locomotor activity of the individual (33). For each of the five treatments there was a matching sham control treatment. Mosquitoes were then placed into a climate chamber (27 °C ±1, 80% humidity and 14:10h light:dark cycle) for 24h before experiments.

### 2.4 Behavioral assay set-up description

In this behavioral assay, we determined which of the *An. gambiae ss* sensory appendages were involved in the non-contact detection of permethrin based on their behavioral response (i.e., takeoff). For this, we used a modified close proximity response assay as in (13) (Fig. 1.A). In brief, the set-up consists of a sample bottle containing a dose of the chemical to be tested which is placed in a water bath allowing the bottle headspace temperature to adjust to 35°C. The bottle’s headspace temperature was measured using a digital thermometer (t110, Testo AG, Germany). The temperature and permethrin concentration were chosen to ensure a positive behavioral response after stimuli exposure [see (13)], as the objective of the experiment was to see whether appendage manipulation would halt the behavioral response. A charcoal-purified continuous airflow was circulated through the sample bottle by means of a stimulus controller (CS-55 v.2, Syntech, Germany). The airflow exiting the bottle was directed by polytetrafluoroethylene (PTFE) tubing towards a single 4- to 8-day-old female mosquito enclosed in a cylindrical cage (⌀10 × L20 cm) covered with mosquito netting. The end of the PTFE tube was positioned at approximatively 2 cm from the individual. Exposure started after the mosquito had been standing still on the netting for at least 5 minutes. The behavior of the mosquito was recorded as a binary variable (i.e., takeoff: yes/no) during a 30s period of exposure to the bottle’s headspace airflow. Each individual was exposed to a single 30s headspace puff.

**Figure 1:**
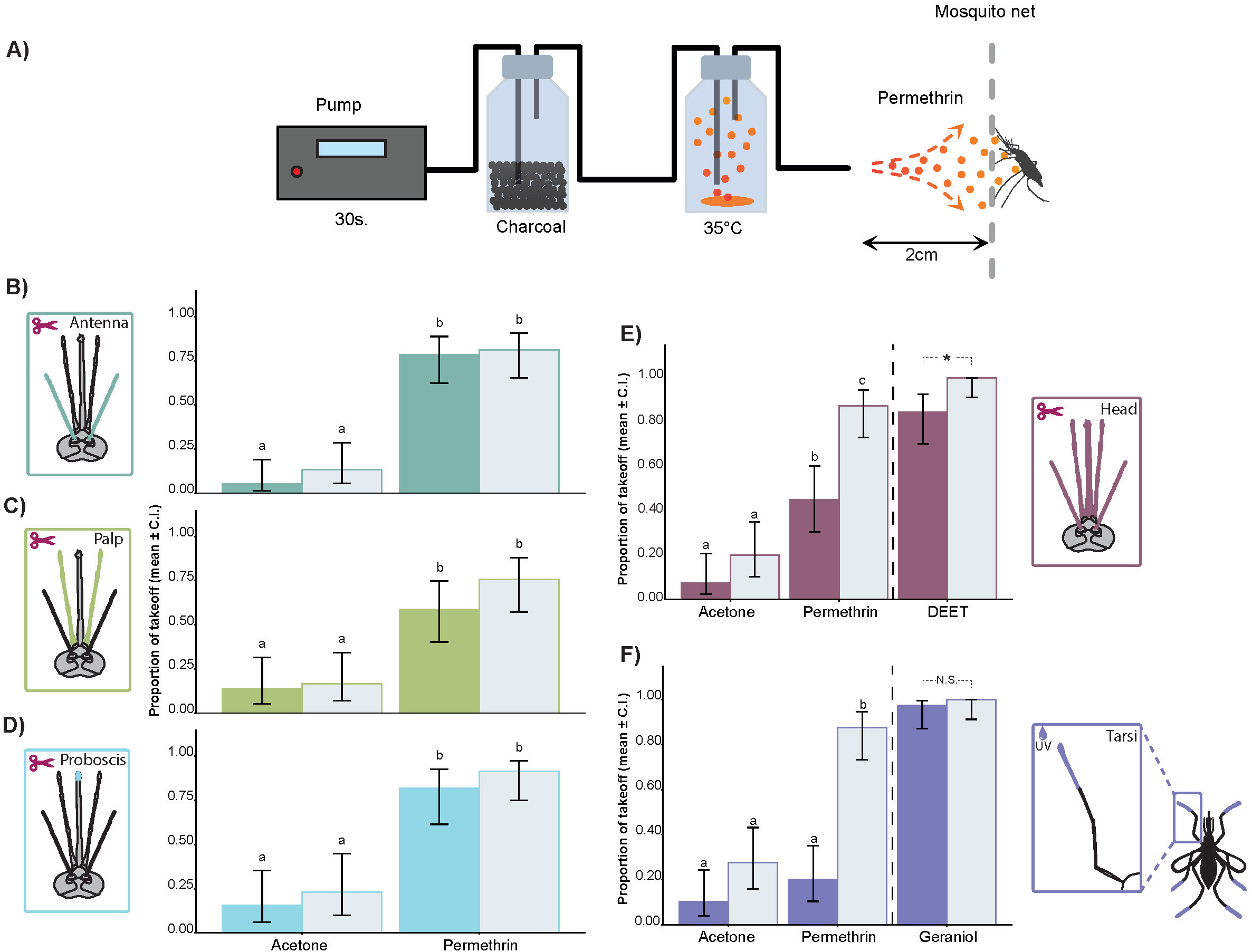
*An. gambiae* takeoff responses after permethrin stimulation depending on the chemosensory appendage manipulated. A) Schematic drawing illustrating the close proximity response assay [see (13) for details].; B) ablated vs non-ablated antennae; C) ablated vs non-ablated palps; D) ablated vs non-ablated proboscis; E) ablated head chemosensory appendages (i.e., simultaneous ablation of antennae, palpi and proboscis) vs non-ablated head chemosensory appendages; F) UV-resin recovered vs non-recovered tarsi. Different letters over bars represent statistically significant differences (see Table S3 for the results of the Tukey’s method for multiple comparisons). * = significant statistical takeoff response between ablated vs non-ablated head chemosensory appendages after DEET stimulation (Fisher exact test for count data: p=0.012). N.S.= not significant differences between resin recovered vs. non-recovered tarsi after geraniol stimulation (Fisher exact test: p=1).

A volume of 1600 μL of a permethrin solution in acetone at 1 mg/mL was used for the test treatment. The same volume of acetone was used as the negative control. A positive control treatment was added for treatments (iv) and (v) to assess whether the takeoff behavior was still observed after appendage manipulation. The sample bottle for the positive control of treatment (iv) contained 40 μL of pure DEET or 40 μL of geraniol solution in acetone at 30% v/v for treatment (v). These compounds have been shown to be detected by other mosquito species via different sensory appendages, i.e., DEET through the tarsi (33) and geraniol through the antennae (34). For each permethrin and control preparations, the solvent was allowed to completely evaporate by sporadically agitating the open bottles for at least 30 minutes under a fume hood at room temperature before the assay.

The number of mosquitoes used in these behavioral assays and the number of replicates for each treatment (insecticide *versus* solvent control) are summarized in Table S2 of the supplementary information.

### 2.5 Electrophysiology: electroantennogram (EAG), electropalpogram (EPG) and electrotarsogram (ETG)

EAG, EPG and ETG responses to test compounds were recorded from individual 4- to 8-day-old *An. gambiae* females. The base of the excised head (for EAGs and EPGs) or the proximal part of the leg (for ETGs) were mounted into glass capillaries (I.D: 0.78 mm, O.D: 1.12 mm), filled with saline Ringer solution and connected to the reference electrode; the distal part of each appendage was also mounted into glass capillaries with saline solution, and connected to the recording electrode. A new appendage preparation was used for every replicate. In total, thirteen replicates (n=13) for EAG, ten (n=10) for EPG and thirteen (n=13) for ETG were performed.

A stimulus controller (CS-55 v.2, Syntech, Germany) was used to deliver charcoal-filtered and humidified air (500 ml/min) continuously over the appendage preparation, by means of silicon tubing connected to a 15 cm long borosilicate glass tube (0.5 cm ID) at the end. The headspace (4.2 ml) of a compound-loaded stimulus pipette was delivered, also through the stimulus controller, via an air puff lasting 0.5 seconds into the continuous air stream. To ensure complete mixing of the test compounds with the continuous air flow, the headspace of the stimuli pipettes was delivered into the continuous air stream through a side port located 10 cm from the end of the borosilicate glass tube.

Stimuli pipettes were loaded by applying 10 µl of the compounds to be tested onto filter paper (0.5 × 2 cm), from which the solvent was allowed to evaporate for about 5 min at room temperature. The filter paper was then inserted inside a glass Pasteur pipette and capped with a 1mL pipette tip. A new set of stimulus pipettes were used for each mosquito preparation. As the headspace airflow in the behavioral experiments was presented to the mosquitoes at 35 °C, we considered it appropriate to keep the same headspace temperature for the electrophysiological experiments. To achieve the chosen temperature of the stimulus pipette headspace, pipettes contrasts between the different were performed using the ‘emmeans’ function of the package ‘emmeans’ (36), adjusted with Tukey HSD to correct for multiple comparisons. Takeoff response differences between manipulated and SHAM individuals when stimulated with the positive controls (DEET and Geraniol) were assessed by Fisher exact test for count data.

For the electrophysiological data, the responses (deflections) in millivolts (mV) were recorded for each appendage in response to the four insecticide concentrations used. Response values were standardized by the response of the same appendage to the negative control stimulus (acetone). These standardized responses were log-transformed for analysis, such that, the transformed variable equals zero when an appendage responded equally to the insecticide and the negative control stimulus. A linear mixed model was fitted to the transformed electrophysiological data using the ‘lmer’ function of the package ‘lme4’ allowing for random effects at the individual level (i.e., accounting for the repeated measures on the same individual). Contrasts were performed to assess whether there was positive electrophysiological response for each of the permethrin doses used. For this, mean response values were compared to zero using the ‘emmeans’ function of the package ‘emmeans’ (36).

## 3 RESULTS

### 3.1 Identification of the chemosensory appendages of *Anopheles gambiae* mosquitoes eliciting a takeoff response during permethrin headspace stimulation

In treatments (i)-(iii), we independently ablated the main sensory appendages of the mosquitoes (i.e., antennae, palpi or proboscis) to decipher which were involved in the non-contact detection of permethrin. We found no statistically significant decrease of the takeoff response when exposed to the permethrin bottle headspace after manipulating any of these sensory appendages (Table 1; Fig. 1B-D).

**Table 1:** Analysis of deviance table based on the proportion of *An. gambiae* taking off after the ablation of the sensory appendage(s) or recovering with resin (tarsi). LR ꭓ^2^=logistic regression model for the effect of the chemical stimuli (acetone vs permethrin), appendage manipulation (ablation/resin recovered vs. SHAM), and their interaction. df= degrees of freedom; P-value (> ꭓ^2^) = probability of obtaining a ꭓ^2^ value greater than the one shown under the null hypothesis. P-value < 0.05 is considered as significant.

| | Variable | LR $\chi^2$ | df | P-value |
| --- | --- | --- | --- | --- |
| Antennae | Chemical stimuli | 82.94 | 1 | **2e-16** |
|  | Appendage manipulation | 0.88 | 1 | 0.3486 |
|  | Interaction | 0.67 | 1 | 0.4139 |
| Palps | Chemical stimuli | 35.23 | 1 | **2.935e-09** |
|  | Appendage manipulation | 1.69 | 1 | 0.1939 |
|  | Interaction | 0.38 | 1 | 0.5372 |
| Proboscis | Chemical stimuli | 48.34 | 1 | **3.592e-12** |
|  | Appendage manipulation | 2.13 | 1 | 0.1447 |
|  | Interaction | 0.18 | 1 | 0.6732 |
| Head sensory appendages | Chemical stimuli | 54.84 | 1 | **1.308e-13** |
|  | Appendage manipulation | 18.64 | 1 | **1.577e-05** |
|  | Interaction | 1.18 | 1 | 0.2766 |
| Tarsi | Chemical stimuli | 27.88 | 1 | **1.292e-07** |
|  | Appendage manipulation | 38.72 | 1 | **4.887e-10** |
|  | Interaction | 5.51 | 1 | **0.01896** |

In treatment (iv), we ablated the three chemosensory appendages simultaneously to test their concomitant implication in the non-contact detection of permethrin. Removing the three appendages significantly reduced the takeoff response of mosquitoes exposed to the permethrin bottle headspace compared to sham control mosquitoes with these appendages (Table 1; Fig. 1E). Yet, the takeoff response of this manipulated group was not completely eliminated as it was significantly greater than the corresponding negative control treatment, i.e., acetone stimulation of mosquitoes with the three head chemosensory appendages obliterated (Table S3, supplementary information). In addition, the positive control (i.e., DEET headspace stimulation) showed mosquitoes were still able to takeoff despite the removal of their head chemosensory appendages (Fig. 1E).

In treatment (v) covering the tarsi with resin significantly reduced the mosquito takeoff response to permethrin headspace stimulation (Table 1; Fig. 1F) to a level similar to the corresponding negative control treatment (i.e., acetone headspace stimulation of mosquitoes with resin covered tarsi) (Table S3, supplementary information). The positive control, 30% geraniol, revealed that despite recovering the tarsi with resin, individuals were still able to takeoff after stimulation (Fig. 1F).

### 3.2 Electrophysiological characterization of permethrin headspace chemosensory detection in *Anopheles gambiae*

The results of the electrophysiological experiments revealed that permethrin headspace elicited antennal (EAG) responses, and the amplitude of this response increased in a dose-dependent manner (ꭓ^2^= 54.36, d.f.=3, Pr <0.0001); Fig. 2B). For antennae, statistically significant EAG responses started from the 0.01% permethrin dose upwards (Table 2). Identifiable electrophysiological responses were absent for the palps at the permethrin concentrations used in this study (Table 2; Fig. 2C), thus no dose-dependent effect was found (ꭓ^2^= 3.82, d.f.=3, Pr =0.28). ETG responses were observed when the mosquito tarsi were stimulated with the permethrin headspace (Fig. 2D), with statistically significant ETG responses starting from the 0.1% permethrin dose upwards (Table 2) in a dose-dependent manner (ꭓ^2^= 67.38, d.f.=3, Pr <0.0001). The amplitude of response to the positive controls for the EAG and EPG were not significantly different when presented before or after permethrin stimulation (Fig. 2B, C), whereas a slight but significant decrease in the response was observed for the ETG (Fig. 2D) after permethrin stimulation.

**Figure 2:**
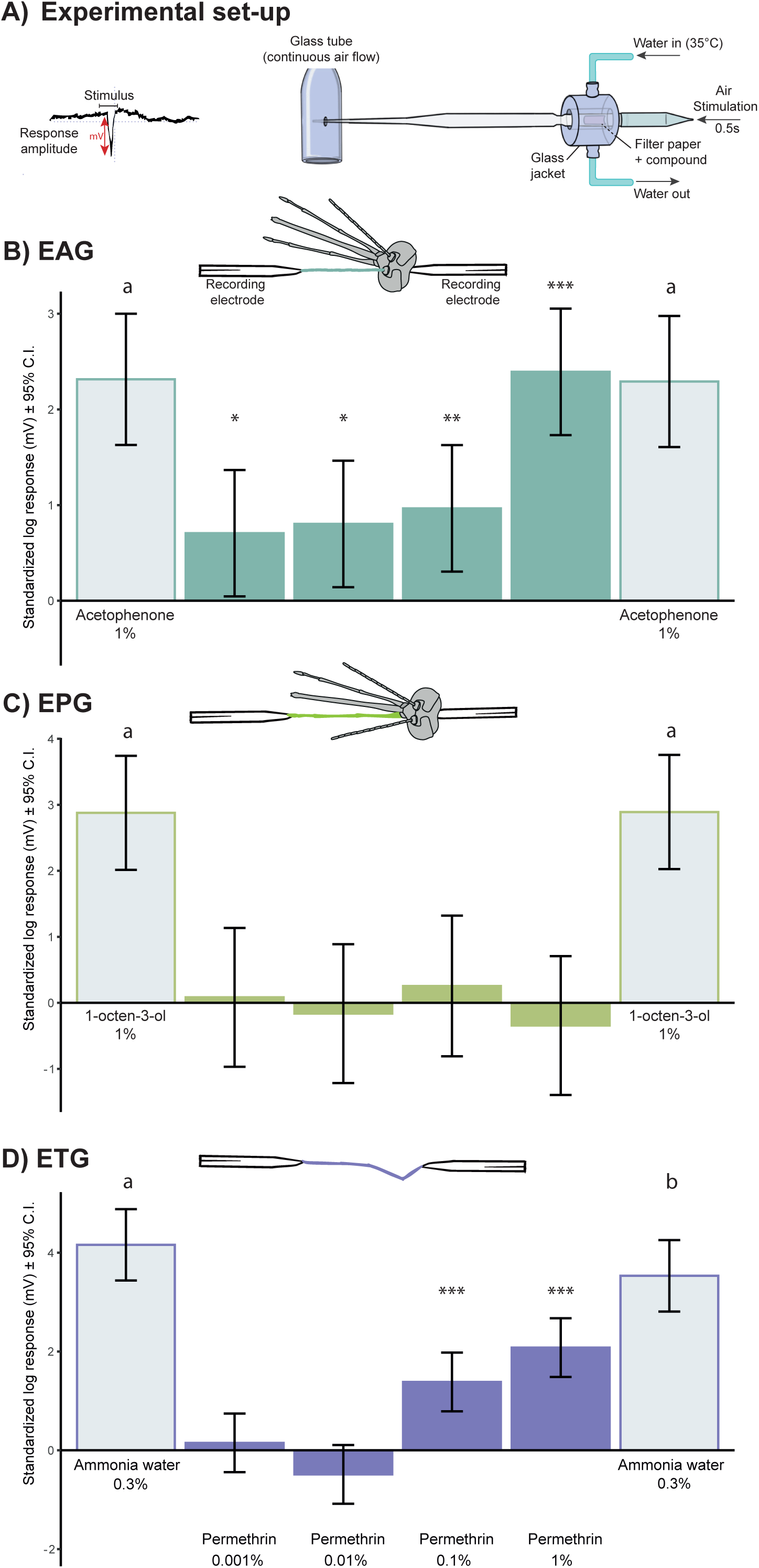
Electrophysiological responses of different *An. gambiae* chemosensory appendages to permethrin. B) Electroantennogram (EAG), (n=13); C) Electropalpogram (EPG), (n=10 mosquitoes); D) Electrotarsogram (ETG), (n=13 mosquitoes). Bars represent the standardized log-response to different doses of permethrin and positive control compounds. Signification of codes: ‘***’ *p<*0.001, ‘**’ *p<*0.01, ‘*’ *p≤*0.05, ‘ ‘ *p>* 0.1. Different letters between positive control compounds indicate significant response differences.

**Table 2:** Contrast table of the electrophysiological responses of different organs depending on the dose of permethrin tested. EAG = Electroantennogram. EPG = Electropalpogram. ETG = Electrotarsogram. S.E. = standard error. LCI= lower confidence interval. UCI = upper confidence interval. P-value < 0.05 is considered as significant.

|  | Dose (%) | EM Mean | S.E. | df | LCI | UCI | t-ratio | P-value |
| --- | --- | --- | --- | --- | --- | --- | --- | --- |
| EAG | 0.001 | 0.71 | 0.34 | 19.6 | 0.01 | 1.41 | 2.09 | **0.049** |
|  | 0.01 | 0.80 | 0.34 | 19.6 | 0.10 | 1.51 | 2.33 | **0.027** |
|  | 0.1 | 0.97 | 0.34 | 19.6 | 0.26 | 1.67 | 2.86 | **0.009** |
|  | 1 | 2.39 | 0.34 | 19.6 | 1.69 | 3.10 | 7.09 | **<0.0001** |
| EPG | 0.001 | 0.08 | 0.54 | 11.9 | -1.09 | 1.25 | 0.15 | 0.879 |
|  | 0.01 | -0.16 | 0.54 | 11.9 | -1.33 | 1.01 | -0.30 | 0.766 |
|  | 0.1 | 0.26 | 0.54 | 12.5 | -0.93 | 1.44 | 0.47 | 0.647 |
|  | 1 | -0.34 | 0.54 | 11.9 | -1.51 | 0.83 | -0.64 | 0.534 |
| ETG | 0.001 | 0.15 | 0.30 | 35.4 | -0.46 | 0.77 | 0.50 | 0.621 |
|  | 0.01 | -0.48 | 0.30 | 35.4 | -1.10 | 0.13 | -1.61 | 0.117 |
|  | 0.1 | 1.38 | 0.30 | 35.4 | 0.77 | 1.99 | 4.57 | **0.0001** |
|  | 1 | 2.08 | 0.30 | 35.4 | 1.46 | 2.69 | 6.86 | **<0.0001** |

## 4 DISCUSSION

This study reveals two chemosensory appendages, tarsi and antennae, involved in the non-contact sensory detection of permethrin for *An. gambiae* mosquitoes. The deflections observed during the electrophysiological recordings on both appendages confirm that both appendages possess the required chemosensory tools to detect permethrin in vapor phase. Nonetheless, the behavioral experiments show the takeoff response of mosquitoes to the dose of permethrin tested in this study was only completely eliminated when tarsi were rendered non-functional at a chemosensory level (i.e., covered in resin), whereas individuals lacking the antennae did not modify their takeoff response compared to the matching control treatment. These results indicate that the behavioral response after exposure to the permethrin headspace is tightly associated to the detection of the molecule by the chemosensory system present in the tarsi.

Tarsal segments are the first to come into contact with a substrate when mosquitoes land, for instance on a flower, on a host or on an insecticide impregnated surface. Tarsal segments are mostly covered with trichoid uniporous gustatory sensilla (29,37). Gustatory sensilla in mosquitoes are usually innervated by the dendrites of 4 chemosensory neurons, which present on their membranes gustatory receptors (GR), Ionotropic receptors (IR) and potentially TRP channels and *pickpocket* receptors (25,30,38). Upon contact, chemical molecules would reach the chemoreceptors by passing through the single pore located at the tip of the sensillum. Hence, these sensors can provide information to the mosquito as to the chemical characteristics of the surface on which it has just landed. In cases where the landing surface contains potentially noxious chemicals, such as insecticides or repellents, mosquitoes usually exhibit a takeoff response and fly away (39). This behavior is commonly referred to as an excito-repellency response in the mosquito literature, and requires direct contact with the chemical impregnated surface. Nonetheless, in this study we observed leg-mediated takeoff responses elicited by permethrin without mosquitoes making direct contact with the impregnated surface. Two hypotheses can be formulated in order to explain these behavioral results. The first hypothesis is that the permethrin being delivered would have deposited and accumulated on the netting tissue of our behavioral set-up, where mosquitoes were standing. In this case, mosquitoes would have detected the insecticide in the same manner as if the surface would have been impregnated with the chemical. The second hypothesis is that mosquito tarsal chemosensory receptors can detect airborne molecules, analogous to the chemosensory process guided by olfactory receptors in the antennae, for instance. The presence of positive responses in our electrophysiological recordings (ETG) are in favor of this second hypothesis. Moreover, a recent study using state-of-the-art single sensillum recordings of gustatory sensilla has shown that dose-dependent neural responses can be elicited before contact with the chemical (40). These results open the question of whether mosquito tarsi, or those of insects in general, serve beyond their taste function and can provide sensory input of airborne molecules at given concentrations.

Permethrin also elicited electrophysiological responses from mosquito antennae (EAG) with a sensitivity 100-fold greater than that of legs (Fig. 2B). This indicates mosquitoes detect this compound through the antennal sensory pathway as well. Yet, ablating the antennae during the behavioral experiments did not suppress the takeoff response during permethrin exposure. This suggests that, although permethrin is detected through the antennal sensory pathway, the information obtained does not elicit a behavioral response from the individual. This raises the question of what the evolutionary advantages would be for *An. gambiae* to possess a seemingly redundant capacity to detect permethrin by different sensory appendages. One potential advantage could be related to learning processes. Mosquitoes, as well as other insects, are able to acquire and retain information in the form of memory (41,42), and several studies have evidenced their olfactory learning capacity (43–45). The antennal detection of permethrin, despite being more sensitive than the tarsal detection, did not elicit any behavioral response in *An. gambiae*. Hence, permethrin could be considered as a neutral stimulus when detected through the antennae at the dose tested in our behavioral experiments. Yet, the antennal sensory input could acquire significance in subsequent exposures to the insecticide if such input is paired with the chemosensory information arriving from the tarsi, which consistently elicits a takeoff response. In a previous study, *Aedes aegypti* and *Culex quinquefasciatus* individuals significantly avoided insecticide impregnated surfaces, including permethrin, after a single previous contact with a non-lethal dose of the insecticide compared to naïve counterparts (46). It is likely that *An. gambiae* individuals would behave in a similar manner as the other two mosquito species when pre-exposed to these molecules, albeit this hypothesis remains to be tested. Incorporating, after a previous non-lethal experience with the molecule, a 100-fold more sensitive chemosensory pathway into the recognition of insecticides by mosquitoes may be highly advantageous in terms of their fitness. Future studies should explore whether mosquitoes can learn to avoid insecticides, particularly in mosquito populations with a high prevalence of physiological insecticide-resistance among individuals.

In the behavioral experiments, removing simultaneously the main chemosensory appendages from the head of mosquitoes (i.e., ablation of antennae, palpi and proboscis) resulted in the partial inhibition of their takeoff response, thus differing from the results obtained in both: i) when the tarsi were occluded, where the takeoff response was consistently halted, and ii) when head chemosensory appendages were individually ablated and the takeoff response was not eliminated. Two hypotheses may explain the results obtained: on one hand, head chemosensory appendages may contribute to ratify, to some extent, the information acquired through the tarsi, as the expected positive takeoff response for having intact tarsi was partially suppressed. On the other hand, a substantial manipulation of chemosensory appendages is presumed to hinder individuals from obtaining most of the environmental chemosensory input and likely alter their behavior (47) for instance in this case to exhibit a takeoff response after stimulation. The positive control treatment proved that mosquitoes were still able to exhibit a takeoff response in the presence of DEET headspace despite the invasive ablation procedure, hence validating the adequacy of our behavioral set-up to answer the question. Nonetheless, DEET and permethrin could have different chemosensory integration pathways, which could explain the differing results obtained with these two molecules. Further studies are needed to precisely characterize how the different chemosensory appendages in the head of Anopheles mosquitoes interact in detecting permethrin.

Finally, the results showing a consistent takeoff response after DEET stimulation suggest that, as well as *Aedes aegypti* (33), *An. gambiae* may also be able to detect this molecule through their tarsi. A previous study with *An. gambiae* showed DEET does not seem to act as a spatial olfactory repellent in this species (34), and suggested that as well as in other species, it may act as a contact repellent. Interestingly, and although using a similar close proximity response assay as in Afify and colleagues, the results of the two experiments are markedly contrasting. This discrepancy is likely to be generated by the way in which stimuli were delivered in the two experiments: i.e., in our study, 100% DEET was heated to 35 °C (within the range of normal skin temperature) and delivered by a continuous airflow, whereas in the study of Afify and colleagues 100% DEET was placed into a pipette tip at room temperature without air delivery. A higher volatilization of the compound due to temperature is expected in our setup, and consequently mosquitoes in our experiment would have received a higher concentration of the compound compared to the study by Afify and colleagues. Thus, DEET’s spatial repellency in *Anopheles* mosquitoes may be correlated to the number of molecules present in the air. Future studies should investigate the concentration thresholds and the role of the different chemosensory pathways for the repellency of Anopheles mosquitoes to repellent compounds.

## 5 CONCLUSION

In this study, we have shown that specific appendages with potentially different chemosensory receptor expression (25), such as the tarsi and the antennae, are able to detect permethrin headspace in *An. gambiae*. Permethrin did not elicit any distinguishable electrophysiological response in the palps at the doses tested in our experiments. The specificity of appendage detection of permethrin and the lack of takeoff response in tarsi-occluded mosquitoes shown in this study, together with the lack of differential takeoff response between insecticide-resistant and insecticide-susceptible strains (13) do not support there being a sensory input mediated by voltage-gated sodium channels agonists (15) in *An. gambiae*. Nonetheless, further functional studies should reveal the role of voltage-gated sodium channels, and other chemosensory-related molecular components such as sensory appendage proteins (48) in eliciting behavioral responses and/or their potential as synergists of insecticide chemosensory-guided behaviors.

## 6 ACKNOWLEDGEMENTS

The authors express their gratitude to Carole Ginibre and Bethsabée Scheid for mosquito rearing and Vectopôle facility management.

