## Supplemental Information for "Permethrin elicits chemoreceptive responses on different *Anopheles gambiae* sensory appendages"

**Table S1:** Detailed information of the chemical products used in this study.

| Compound | IUPAC Compound Name | Product number | Brand | CAS number | Purity |
| --- | --- | --- | --- | --- | --- |
| Permethrin | 3-Phenoxybenzyl 3-(2,2-dichlorovinyl)-2,2-dimethylcyclopropanecarboxylate | 45614 | Sigma-Aldrich | 52645-53-1 | 95% |
| DEET | N, N-Diethyl-3-methylbenzamide | 100951 | Sigma-Aldrich | 134-62-3 | 97% |
| Geraniol | (2E)-3,7-Dimethyl-2,6-octadien-1-ol | 48798 | Sigma-Aldrich | 106-24-1 | ≥ 98.5% |
| Acetophenone | 1-Phenylethanone | 42163 | Sigma-Aldrich | 98-86-2 | 99% |
| 1-octen-3-ol | 1-octen-3-ol | 05284 | Sigma-Aldrich | 3391-86-4 | 98% |
| Ammonia water | Azanium; hydroxide | 221228 | Sigma-Aldrich | 1336-21-6 | 30% in water |
| Acetone | 2-Propanone | 650501 | Sigma-Aldrich | 67-64-1 | ≥ 99.9% |

**Table S2:** Sample sizes of mosquitoes used in behavioral tests. N= total number of mosquitoes combining all replicates for a given treatment.

| **Sensory organ** | **Treatment** | **Acetone (control)** | | **Permethrin** | | **DEET** | | **Geraniol** | |
| --- | --- | --- | --- | --- | --- | --- | --- | --- | --- |
|  |  | N | Replicates | N | Replicates | N | Replicates | N | Replicates |
| Antennae | SHAM | 37 | 4 | 37 | 4 | _ | | _ | |
|  | Ablated | 37 | 4 | 37 | 4 | _ | | _ | |
| Palpi | SHAM | 30 | 3 | 30 | 3 | _ | | _ | |
|  | Ablated | 30 | 3 | 30 | 3 | _ | | _ | |
| Proboscis | SHAM | 40 | 4 | 40 | 4 | _ | | _ | |
|  | Ablated | 40 | 4 | 40 | 4 | _ | | _ | |
| Head chemosensory organs | SHAM | 40 | 4 | 40 | 4 | 40 | 4 | _ | |
|  | Ablated | 40 | 4 | 40 | 4 | 40 | 4 |  | |
| Tarsi | SHAM | 40 | 4 | 40 | 4 | _ | | 40 | 4 |
|  | Resin covered | 40 | 4 | 40 | 4 | _ | | 40 | 4 |

**Table S3:** Pairwise contrasts (Tukey HSD method) of the takeoff response from *An. gambiae* mosquitoes after permethrin or acetone stimulation. T1 vs. T2 = treatments being contrasted. S.E. = standard error. LCL= lower confidence limit. UCL = upper confidence limit. P-value < 0.05 is considered as significant.

| **Organ**  **manipulated** | **Contrast** | | | | | **Odds Ratio** | **S.E.** | **LCL** | **UCL** | **Z-ratio** | **P-value** |
| --- | --- | --- | --- | --- | --- | --- | --- | --- | --- | --- | --- |
|  | **T1** | | ***vs.*** | **T2** | |  |  |  |  |  |  |
|  | **Stimulus** | **Manipulation** |  | **Stimulus** | **Manipulation** |  |  |  |  |  |  |
| Antennae | Acetone | SHAM | *vs.* | Permethrin | SHAM | 0.04 | 0.02 | 0.01 | 0.19 | -5.19 | **<.0001** |
|  | Acetone | SHAM | *vs.* | Acetone | Ablation | 2.73 | 2.38 | 0.29 | 25.67 | 1.15 | 0.6559 |
|  | Acetone | SHAM | *vs.* | Permethrin | Ablation | 0.04 | 0.03 | 0.01 | 0.22 | -5.03 | **<.0001** |
|  | Permethrin | SHAM | *vs.* | Acetone | Ablation | 75.00 | 63.00 | 8.68 | 648.17 | 5.14 | **<.0001** |
|  | Permethrin | SHAM | *vs.* | Permethrin | Ablation | 1.18 | 0.69 | 0.27 | 5.24 | 0.29 | 0.9916 |
|  | Acetone | Ablation | *vs.* | Permethrin | Ablation | 0.02 | 0.01 | 0.01 | 0.13 | -5.00 | **<.0001** |
| Palpi | Acetone | SHAM | *vs.* | Permethrin | SHAM | 0.06 | 0.04 | 0.01 | 0.34 | -4.21 | **0.0002** |
|  | Acetone | SHAM | *vs.* | Acetone | Ablation | 1.25 | 0.91 | 0.19 | 8.11 | 0.31 | 0.9900 |
|  | Acetone | SHAM | *vs.* | Permethrin | Ablation | 0.14 | 0.09 | 0.03 | 0.69 | -3.17 | **0.0084** |
|  | Permethrin | SHAM | *vs.* | Acetone | Ablation | 19.64 | 13.60 | 3.32 | 116.10 | 4.31 | **0.0001** |
|  | Permethrin | SHAM | *vs.* | Permethrin | Ablation | 2.22 | 1.28 | 0.51 | 9.72 | 1.39 | 0.5081 |
|  | Acetone | Ablation | *vs.* | Permethrin | Ablation | 0.11 | 0.07 | 0.02 | 0.61 | -3.32 | **0.0050** |
| Proboscis | Acetone | SHAM | *vs.* | Permethrin | SHAM | 0.03 | 0.02 | 0.01 | 0.17 | -5.17 | **<.0001** |
|  | Acetone | SHAM | *vs.* | Acetone | Ablation | 1.62 | 0.93 | 0.37 | 7.04 | 0.84 | 0.8338 |
|  | Acetone | SHAM | *vs.* | Permethrin | Ablation | 0.07 | 0.04 | 0.02 | 0.30 | -4.65 | **<.0001** |
|  | Permethrin | SHAM | *vs.* | Acetone | Ablation | 57.40 | 41.60 | 8.92 | 369.36 | 5.59 | **<.0001** |
|  | Permethrin | SHAM | *vs.* | Permethrin | Ablation | 2.35 | 1.59 | 0.41 | 13.39 | 1.26 | 0.5860 |
|  | Acetone | Ablation | *vs.* | Permethrin | Ablation | 0.04 | 0.03 | 0.01 | 0.20 | -5.13 | **<.0001** |
| Head Chemosensory Organs | Acetone | SHAM | *vs.* | Permethrin | SHAM | 0.04 | 0.02 | 0.01 | 0.18 | 5.37 | **<.0001** |
|  | Acetone | SHAM | *vs.* | Acetone | Ablation | 3.08 | 2.22 | 0.49 | 19.54 | 1.57 | 0.3978 |
|  | Acetone | SHAM | *vs.* | Permethrin | Ablation | 0.31 | 0.16 | 0.08 | 1.13 | -2.34 | 0.0897 |
|  | Permethrin | SHAM | *vs.* | Acetone | Ablation | 86.33 | 66.30 | 12.02 | 620.01 | 5.81 | **<.0001** |
|  | Permethrin | SHAM | *vs.* | Permethrin | Ablation | 8.56 | 4.91 | 1.96 | 37.39 | 3.74 | **0.0011** |
|  | Acetone | Ablation | *vs.* | Permethrin | Ablation | 0.10 | 0.07 | 0.02 | 0.567 | -3.40 | **0.0037** |
| Tarsi | Acetone | SHAM | *vs.* | Permethrin | SHAM | 0.05 | 0.03 | 0.01 | 0.25 | -4.90 | **<.0001** |
|  | Acetone | SHAM | *vs.* | Acetone | Resin | 3.32 | 2.11 | 0.65 | 16.99 | 1.89 | 0.2333 |
|  | Acetone | SHAM | *vs.* | Permethrin | Resin | 1.52 | 0.81 | 0.39 | 5.93 | 0.79 | 0.8611 |
|  | Permethrin | SHAM | *vs.* | Acetone | Resin | 61.25 | 43.60 | 9.83 | 381.61 | 5.78 | **<.0001** |
|  | Permethrin | SHAM | *vs.* | Permethrin | Resin | 28.00 | 17.40 | 5.69 | 137.81 | 5.37 | **<.0001** |
|  | Acetone | Resin | *vs.* | Permethrin | Resin | 0.46 | 0.30 | 0.08 | 2.49 | -1.19 | 0.635 |
